# Deciphering the role of the Pancreatic Secretome in Covid-19 associated Multi-Organ Dysfunctions

**DOI:** 10.1101/2021.09.22.461447

**Authors:** Ekta Pathak, Rajeev Mishra

**Author notes:** To whom correspondence should be addressed: **Dr. Ekta Pathak,** Ph.D. in Bioinformatics, Banaras Hindu University, Bioinformatics Scientist (Freelance), **Dr. Rajeev Mishra,** Associate Professor & Coordinator, Bioinformatics, MMV, Institute of Science, Banaras Hindu University, Varanasi-221005, India.

## Abstract

Emerging evidence indicates an intricate relationship between the SARS-CoV-2 infection and Multi-Organ Dysfunctions (MODs). Here, we have investigated the role of the Secretome of the SARS-CoV-2 infected pancreas and mechanistically linked it with the multi-organ dysfunction using the scRNA-seq analysis. We found that acinar-specific *PRSS2, REG3A, REG1A, SPINK1*, and ductal-specific *SPP1, MMP7* genes are upregulated in alpha, beta, delta, and mesenchyme cells. Using extensive documented experimental evidence, we validated the association of upregulated pancreatic Secretome with coagulation cascade, complement activation, renin angiotensinogen system dysregulation, endothelial cell injury and thrombosis, immune system dysregulation, and fibrosis. Our finding suggests the influence of upregulated Secretome on multi-organ systems such as Nervous, Cardiovascular, Immune, Digestive, and Urogenital systems. In addition, we report that the secretory proteins IL1B, AGT, ALB, SPP1, CRP, SERPINA1, C3, TFRC, TNFSF10, and MIF are associated with diverse diseases. Thus, suggest the role of the pancreatic Secretome in SARS-CoV-2 associated MODs.

## Introduction

The ongoing pandemic Coronavirus disease 2019 (COVID-19) caused by severe acute respiratory syndrome coronavirus 2 (SARS-CoV-2) encompass a myriad of pathologies(Zhu et al., 2020). The SARS-CoV-2 infection affects the lungs, heart, kidney, intestine, olfactory epithelia, liver, and pancreas and brings forward multi-organ dysfunctions (MODs) in many patients(Chen et al., 2020; Fang et al., 2020; Giacomelli et al., 2020; Lamers et al., 2020; Peiris et al., 2021; Puelles et al., 2020; Yang et al., 2020).

SARS-CoV-2 uses the ACE-2 receptor to enter the host cells and cause pancreatic injury (Liu et al., 2020; Zhou et al., 2020). Acute pancreatitis (AP) initiates in the pancreas in response to an inflammatory event, leading to deleterious local and systemic effects (Götzinger et al., 2003) and eventually leads to multi-organ damage and dysfunction(Bhatia, 2009). There are cases of pancreatitis associated without respiratory symptoms (Kandasamy, 2020; Lakshmanan and Malik, 2020) and after the clearance of SARS-CoV-2 in the lungs(Zhao et al., 2020) of the COVID-19 patients. While the detailed mechanisms of SARS-CoV-2 induced acute pancreatitis are still under investigation(AlHarmi et al., 2021; de-Madaria et al., 2021; Ramos-Casals et al., 2021), the AP pathogenesis is commonly attributed to trypsin activation and intracellular signaling(Frossard, 2001), the release of proteolytic enzymes such as amylase and lipase(Tauseef et al., 2021), reactive oxygen species (ROS) (Tsuji et al., 1994), inflammatory elements, and release of other mediators into the blood, collectively leading to activation of the systemic inflammatory response(Bruen et al., 2021; Vege and Chari, 2021).

Many aspects of SARS-CoV-2 induced organ damage have been investigated (Cardona et al., 2020; Fajgenbaum and June, 2020; Moore and June, 2020; Ronco and Reis, 2020; Sultan et al., 2020; Veras et al., 2020). However, the involvement of specific pathways, such as those centered on pancreatic infection of SARS-CoV-2, needs to be investigated. Here, to mechanistically link the multi-organ dysfunction with COVID-19 infected pancreas, we have investigated the role of upregulated pancreatic secretory proteins (pancreatic Secretome) in COVID-19 associated MODs using scRNA-seq data of ex vivo SARS-CoV-2 infected human pancreas. Using documented experimental evidence, we validated that upregulated pancreatic Secretome is associated with coagulation cascade, complement activation, renin angiotensinogen system dysregulation, endothelial cell injury and thrombosis, immune system dysregulation, and fibrosis. Furthermore, our finding suggests the influence of upregulated Secretome on multi-organ systems such as Nervous, Cardiovascular, Immune, Digestive, And Urogenital systems. In addition, we report that the secretory proteins IL1B, AGT, ALB, SPP1, CRP, SERPINA1, C3, TFRC, TNFSF10, and MIF are associated with diverse diseases. Thus, suggest the role of the upregulated pancreatic Secretome in MODs.

## Results and discussion

Recent reports have indicated the damage in the gastrointestinal mucosal barrier in COVID-19 patients (Massironi et al., 2020; Sharma and Riva, 2020; Vanella et al., 2021). The damage in the mucosal barrier may cause the digestive enzymes of the exocrine pancreas to enter into the systemic circulation (Altshuler et al., 2016). As a result, the pancreatic secretory proteins may reach other body organs, thus, being involved in multi-organ dysfunction. Therefore, we have investigated the role of pancreatic secretions in multi-organ dysfunction using scRNA-seq analysis of SARS-CoV-2 infected pancreas.

### Identification and analysis of Pancreatic cell-type-specific differentially expressed Secretome

We downloaded scRNA-seq data for Mock-infected and SARS-CoV-2 infected pancreas from Gene Expression Omnibus under accession code GSE159556; (Tang et al., 2021). The clustering analysis of the scRNA-seq data showed 45 different clusters. These clusters were merged to form nine clusters of different cell types, i.e., acinar cells, ductal cells, alpha cells, beta cells, delta cells, PP cells, endothelial cells, mesenchyme cells, and immune cells (Fig.1 A). The cell type identification was based on marker genes *PRSS2* (acinar cells), *KRT19* (ductal cells), *GCG* (alpha cells), *INS* (beta cells), *COL1A1* (mesenchyme cells), *PPY* (PP cells), *SST* (delta cells), *ESAM* (endothelial cells), and *LAPTM5* (immune cells) (Fig.1 B), using reported literature (Muraro et al., 2016).

**Fig.1.**
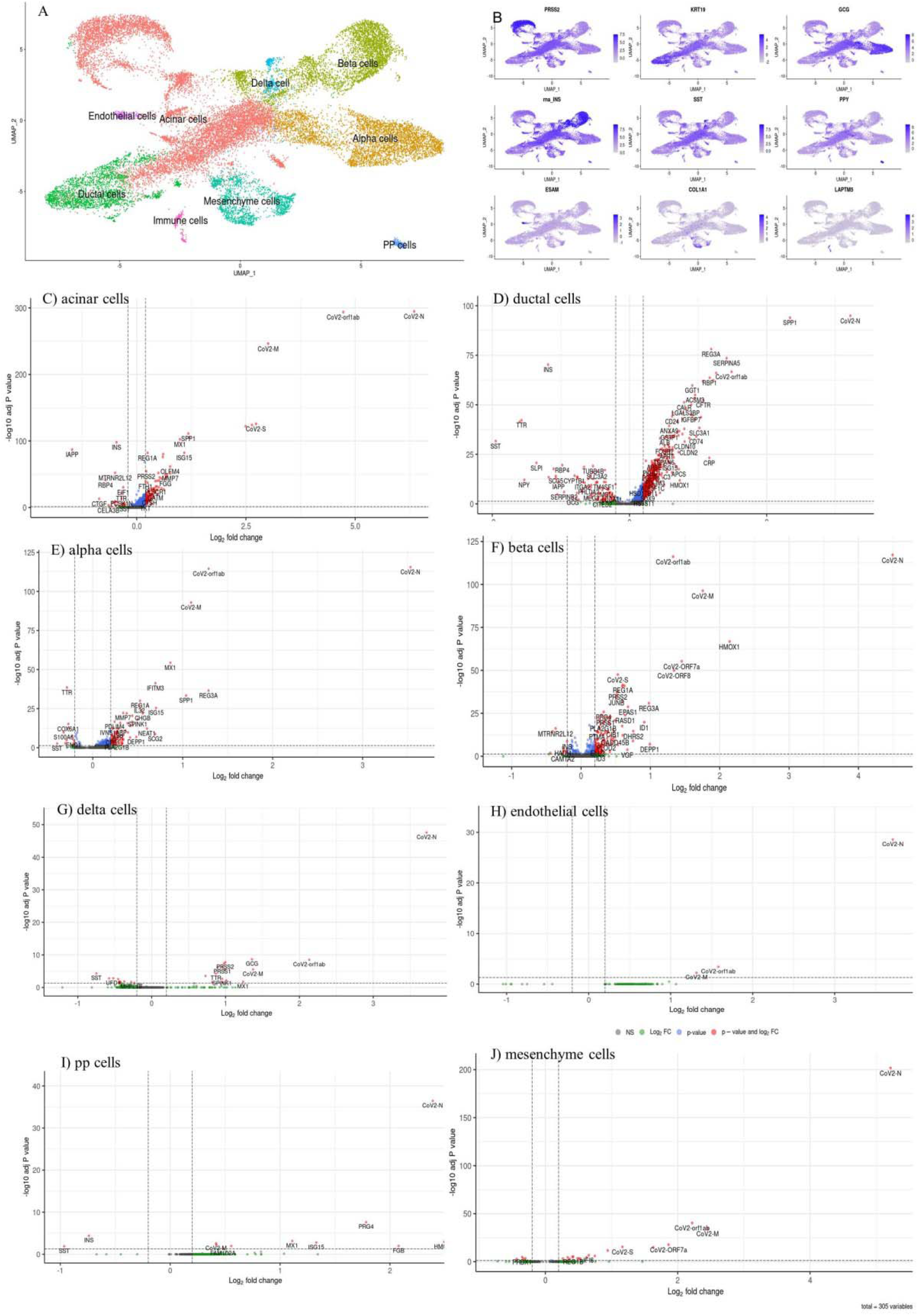
Cell type and DEGs identification. A) UMAP of pancreatic cells showing cell types. B) UMAP of cell marker genes PRSS2(acinar cells), KRT19(ductal cells), GCG(alpha cells), INS(beta cells), COL1A1(mesenchyme cells), PPY(PP cells), SST(delta cells), ESAM(endothelial cells), and LAPTM5(immune cells). Volcano plot showing differentially expressed genes in C) acinar cells, D)ductal cells, E) alpha cells, F) beta cells, G) delta cells, H) PP cells, I)endothelial cells, J)mesenchyme cells

We have identified that *CoV2-N, CoV2-orf1ab, CoV2-M, CoV2-S, CoV2-ORF7a, CoV2-ORF8* viral genes are expressed across all cell types in the single-cell expression analysis (Fig.1 C-J). In addition, we have identified cell type-specific differentially expressed genes: 149 in acinar, 631 in ductal, 107 in alpha, 151 in beta, 28 in delta, 3 in endothelial, 11 in pp cells, and 22 genes in mesenchyme cells. The upregulated genes are 125 genes in acinar, 538 in ductal, 94 in alpha, 139 in beta, 16 in delta, 3 in endothelial, 9 in pp cells, and 18 in mesenchyme cells. A superset of 712 genes is found to be upregulated **(Table S1)**. In addition, we found that *SPINK1, OLFM4, ISG15, REG1A, SPP1, REG3A, MMP7, ALB, IL32, PRSS2, REG1B* genes are upregulated in 4 or more cell types in COVID-19 infection **(Table S1)**. Noticeably, we observed upregulation of acinar-specific genes *PRSS2, REG3A, REG1A, SPINK1*, and ductal-specific genes *SPP1, MMP7* in alpha, beta, delta, and mesenchyme cells also. However, the expression of marker gene GCG does not change significantly in alpha cells. Also, INS expression in the beta cell is downregulated in the COVID-19 condition. Our DEG analysis indicates that COVID -19 infections shift the expression profile of pancreatic endocrine cells to acinar and ductal cell-specific profiles (Fig.1 C-J). Thus, resulting in amplified expression of the acinar and ductal specific genes.

Furthermore, 712 upregulated genes were subjected to identification of the secretory proteins using The Human Protein Atlas (Fig.2A). We identified secretory proteins: 34 in acinar, 65 in ductal, 26 in alpha, 28 in beta, 10 in delta, 3 in pp cells, and 6 in mesenchyme cells. Taken together, we found 102 upregulated pancreatic secretory proteins (pancreatic Secretome). The upregulated Secretome was used to construct the protein-protein interaction network using the string database (Fig.2B). We found ALB, IL1B, SERPINA1, CRP, CD44, TTR, CTSB, VTN, SPP1, C3, MMP7, and AGT proteins to be influential using network topological parameters degree and closeness centrality **(Table S2)**.

**Fig.2.**
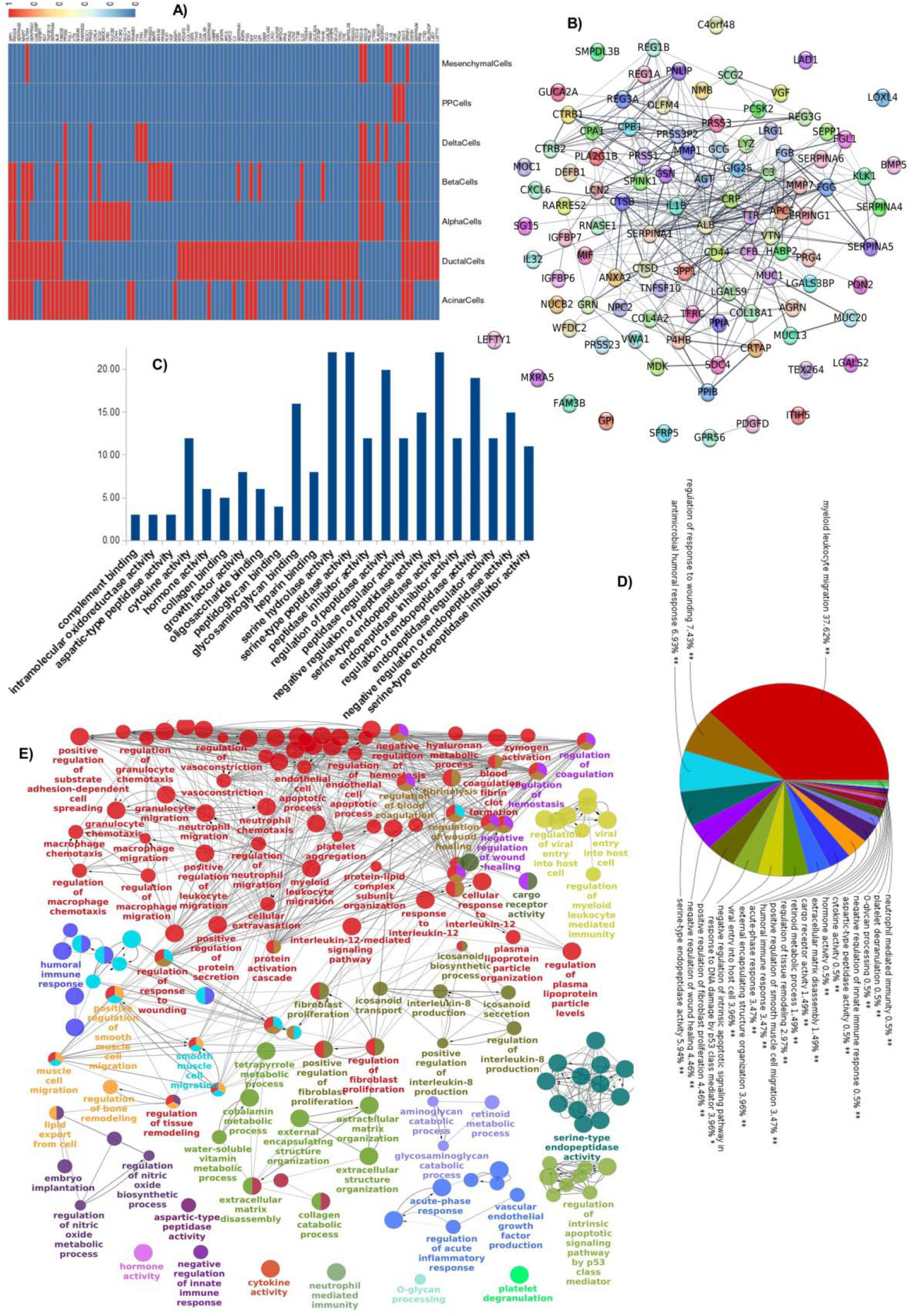
Secretome of SARS-CoV-2 infected Pancreatic cells. A) Heatmap showing up-regulated secretory genes of Covid-19 infected pancreatic cells. Upregulated genes are shown in red color. The blue color indicates that genes are not upregulated. B) Protein-Protein Interaction network of the Secretome of Covid-19 infected pancreatic cells. Enrichment analysis of Secretome: C) molecular function, D) representative biological processes, and E) network of biological processes

### Role of upregulated Pancreatic Secretome

We investigated the role of upregulated *PRSS2, REG3A, REG1A, SPINK1 SPP1, MMP7, OLFM4, ISG15, ALB, IL32, REG1B, AGT, IL1B, SERPINA, CRP, CD44, VTN, TTR*, and *CTSB* genes using the available experimental evidence. The upregulation of PRSS1 and PRSS2 is a characteristic of pancreatitis that causes increased intra-pancreatic trypsin activity resulting in pancreatic damage(Le Maréchal et al., 2006) (Masson et al., 2008). PRSS1 and PRSS2 encode serine protease trypsin that can cleave complement components C3 into C3a and C3b, and C5 into C5a and C5b. C3a and C5a are known to mediate inflammation(Huber-Lang et al., 2018). C3 has a pivotal role in the activation of the complement system. C3 deposition occurs around damaged acinar cells in pancreatitis(Seelig et al., 1978). It induces neutrophil infiltration and neutrophil extracellular traps formation. Neutrophil infiltration is associated with severe acute pancreatitis tissue damage (Linders et al., 2020) (Castanheira and Kubes, 2019). Activated trypsin leads to pancreatic injury and bleeding. Sha. et al. 2009, have reported trypsin to be responsible for multiple organ injuries. It reaches other organs through venous flow circulation.(Sha et al., 2009) Similarly, SPINK1 is overexpressed in pancreatitis, and the elevation is associated with the disease severity(Ohmuraya and Yamamura, 2011). REG1A and REG3A have enhanced expression during pancreatitis. REG1A and REG1B are involved in the regeneration of islet cells and diabetogenesis. REG3A is involved in cell proliferation and has antibacterial activity(Chen et al., 2019). SPP1(Osteopontin) is an extracellular structural protein that binds hydroxyapatite. It is involved in efficient T-helper 1 cell immune responses and enhances mast cell responses to antigen(Nagasaka et al., 2008). SPP1 is a cytokine that upregulates the expression of IL-12 and IFN-γ. IL-12 also induces T-helper 1 cell differentiation and secretion of IFN-γ (Tjan et al., 2021). IFN-γ plays an essential role in defense against virus and activates T cells cytokine production. But, consistently increased IFN-γ level aggravates the systemic inflammation, leading to tissue damage and organ failure(Gadotti et al., 2020). MMP7 degrades casein, gelatins, fibronectin and activates procollagenase. Azzaq Belaaouaj, Abderr, et al. 2000, showed that MMP7 along with MMP1, MMP9, and MMP12 can create a thrombosis-prone environment in atherosclerotic plaques and alter the coagulation pathway in inflammatory diseases(azzaq Belaaouaj et al., 2000). ALB is the main plasma protein and regulates the colloidal osmotic pressure of blood(Sugio et al., 1999). IL32 is a cytokine that induces cytokines such as TNF-α and IL6 and chemokines IL8 and CXCL2. In addition, it activates the signal pathways of NF-kappa-B and p38 MAPK(Kim et al., 2005). ISG15 induces the production of IFN-gamma, ubiquitination of newly-synthesized proteins. It helps in the proliferation of natural killer cells and is a chemotactic factor for neutrophils. It inhibits viral replication and regulates the host’s damage and repair response (Perng and Lenschow, 2018). OLFM4 is a glycoprotein that assists in cell adhesion and is an antiapoptotic factor promoting tumor growth(Gersemann et al., 2012). AGT, a part of the renin-angiotensin system (RAS), regulates blood pressure. Its inhibition reduces atherosclerosis and kidney dysfunction in polycystic kidney disease(Lu et al., 2016). IL-1β is a pro-inflammatory cytokine(Dinarello, 1996). It induces T and B-cell activation, cytokine and antibody production, neutrophil infiltration, and activation(Nakae et al., 2001; Tominaga et al., 2000). It also induces prostaglandin synthesis, fibroblast proliferation, vascular endothelial growth factor (VEGF) production (Fiebich et al., 2000; Nakahara et al., 2003; Siwik et al., 2000). SERPINA1 is serine proteases inhibitor and is reported as a potential prognostic marker of COVID-19(Dutta and Goswami, 2021). CRP is involved in inflammation and helps in complement binding to invaders and apoptotic cells. It helps in opsonin-mediated phagocytosis, production of IL1B, IL6 and TNF-α, and reduces the nitric oxide production(Sproston and Ashworth, 2018). CD44 is a cellular adhesion molecule for extra cellular matrix (ECM) component, hyaluronic acid(Aruffo et al., 1990). VTN an adhesive glycoprotein present in serum and ECM. It repairs and remodels ECM in different tissues after trauma(Leavesley et al., 2013). TTR transports thyroxin and retinol-retinol binding complex to brain and other body parts and induces oxidative stress in endoplasmic stress(Sharma et al., 2019; Teixeira et al., 2006).CTSB is involved in extracellular matrix degradation (Porter et al., 2013). Taken together, PRSS2, REG3A, REG1A, SPINK1 SPP1, MMP7, OLFM4, ISG15, ALB, IL32, and REG1B, AGT, IL1B, SERPINA, CRP, CD44, VTN, TTR and CTSB are involved in complement and coagulation cascade, extra-cellular matrix assembly, fluid balance, and immune response and may lead to sepsis.

### Enrichment analysis of Pancreatic Secretome: GO, Biological pathway, Disease phenotypes

The upregulated Secretome was further analyzed for the GO terms, Biological pathways, Disease phenotypes. We identified functionally grouped GO and pathways using ClueGO analysis (Bindea et al., 2009). We found that serine-type peptidase activity, endopeptidase activity, glycosaminoglycan binding, and cytokine activity were among the top enriched molecular functions (Fig.2 C). We noted that the top enriched biological processes were related to myeloid leukocyte migration (37.62%), regulation of response to wounding (7.43%), antimicrobial humoral response (6.93%), serine-type endopeptidase activity (5.94%), and positive regulation of fibroblast proliferation (4.46%) (Fig.2 D). We found that the biological process of metabolism of tetrapyrrole, cobalamin, hyaluronan and retinoid, and catabolism of collagen, aminoglycan, and glycosaminoglycan were also enriched. We revealed that Myeloid leukocyte migration was associated with neutrophil-mediated immunity, neutrophil chemotaxis, regulation of macrophage migration, positive regulation of protein secretion, endothelial cell apoptotic process, vascular endothelial growth factor production, interleukin-12-mediated signaling pathway, vasoconstriction, zymogen activation, platelet aggregation, and regulation of coagulation. The biological function of fibroblast proliferation was functionally linked to eicosanoid secretion and interleukin-8 production (Fig.2 E). Using the documented experimental evidence, we corroborated the role of enriched biological processes and molecular functions and their implications in MODs. The Endothelial cells (ECs) regulate the coagulation cascade. EC activation and dysfunction are reported in COVID-19 patients (Bermejo-Martin et al., 2020). It interferes with vascular integrity and leads to EC apoptosis activating clotting cascade(Sturtzel and Pathology, 2017). Platelets bind to Cell adhesion molecule (CAM) displayed by activated EC. Platelets secreted Vascular endothelial growth factors (VEGF) induce tissue factor and matrix metalloproteinase production in endothelial cells. This leads to thrombus formation and degradation of the underlying basement membrane, which causes vascular permeability(Zucker et al., 1998). Clinical study shows elevated levels of VEGF in Covid-19 patients(Huang et al., 2020). High levels of VEGF lead to plasma extravasation, edema, and increased tissue hypoxia.VEGF is also involved in atherosclerosis(Yang et al., 2003). As a result of increased endothelial permeability, neutrophil migration occurs (Lehman et al., 2006).In COVID-19, over-activation of neutrophils in response to infection leads to excessive reactive oxygen species (ROS) production. The ROS damage tetrapyrrole rings like heme of hemoglobin and nitric oxide synthase (NOS) and the corrin ring of vitamin B12. The destruction of hemoglobin leads to hypoxia and protein aggregation. Destruction of NOS leads to deficiency of nitric oxide (NO) and ultimately to vasoconstriction. The destruction of the corrin ring results in vitamin B12 deficiency[74], leading to oxidative stress, hypercoagulation, and vasoconstriction (Shakoor et al., 2021). Low levels of NO, Oxygen, and Vitamin B12 deficiency are reported in COVID-19 patients(Siwik and Colucci, 2004). ROS also increases the matrix metalloproteinase (MMP) expression, as evident from our analysis. MMPs are involved in extracellular matrix organization and disassembly, which increases the production of chemokines and cytokines(Siwik and Colucci, 2004). The high molecular weight glycosaminoglycan polymer, hyaluronan (HMW-HA), has anti-inflammatory properties. HMW-HA, in acute inflammation, binds with fibrin and fibrinogen, which leads to increased clot formation. Also, the HMW-HA is broken down into low molecular weight hyaluronan(LMW-HA), and oligo-HA by neutrophils produced ROS. LMW-HA increases the vascular permeability, and both oligo-HA and LMW-HA act as Damage-associated molecular patterns (DAMPs). This leads to aggravated cytokine storms (Ontong and Prachayasittikul, 2021). High levels of hyaluronans are reported in critical COVID-19 patients(Ding et al., 2020). Deficiency of retinol and retinoic acids occurs in SARS-CoV-2 infection due to increased catabolic process that leads to retinoid signaling defect. It causes excessive cytokine secretion, leading to systemic effects and MOD(Sarohan et al., 2021). Eicosanoids are arachidonic acid-derived chemicals and are pro-inflammatory (Hammock et al., 2020). They are involved in physiological processes such as fever, allergy, pain(Serhan, 2014). Eicosanoids are dramatically upregulated in nonsurvivors of sepsis-induced multi-organ dysfunction(Wang et al., 2020). An increased prostaglandins (eicosanoids) level contributes to the cytokine storm (Hosoi et al., 2013).

The pathway enrichment analysis of pancreatic Secretome revealed that the biological pathways were associated with the Pancreatic secretion, RAS and bradykinin pathways in COVID-19, complement and coagulation cascades, IL-17 signaling pathway, ECM-receptor interaction, Protein digestion, and absorption, Type II interferon signaling (IFNG), Vitamin B12 and Folate Metabolism, Lung fibrosis, Hepatitis C and Hepatocellular Carcinoma, Interleukin-12 family signaling, Platelet activation, signaling and aggregation, Gene and protein expression by JAK-STAT signaling (Fig.3 A, B, C).

**Fig.3.**
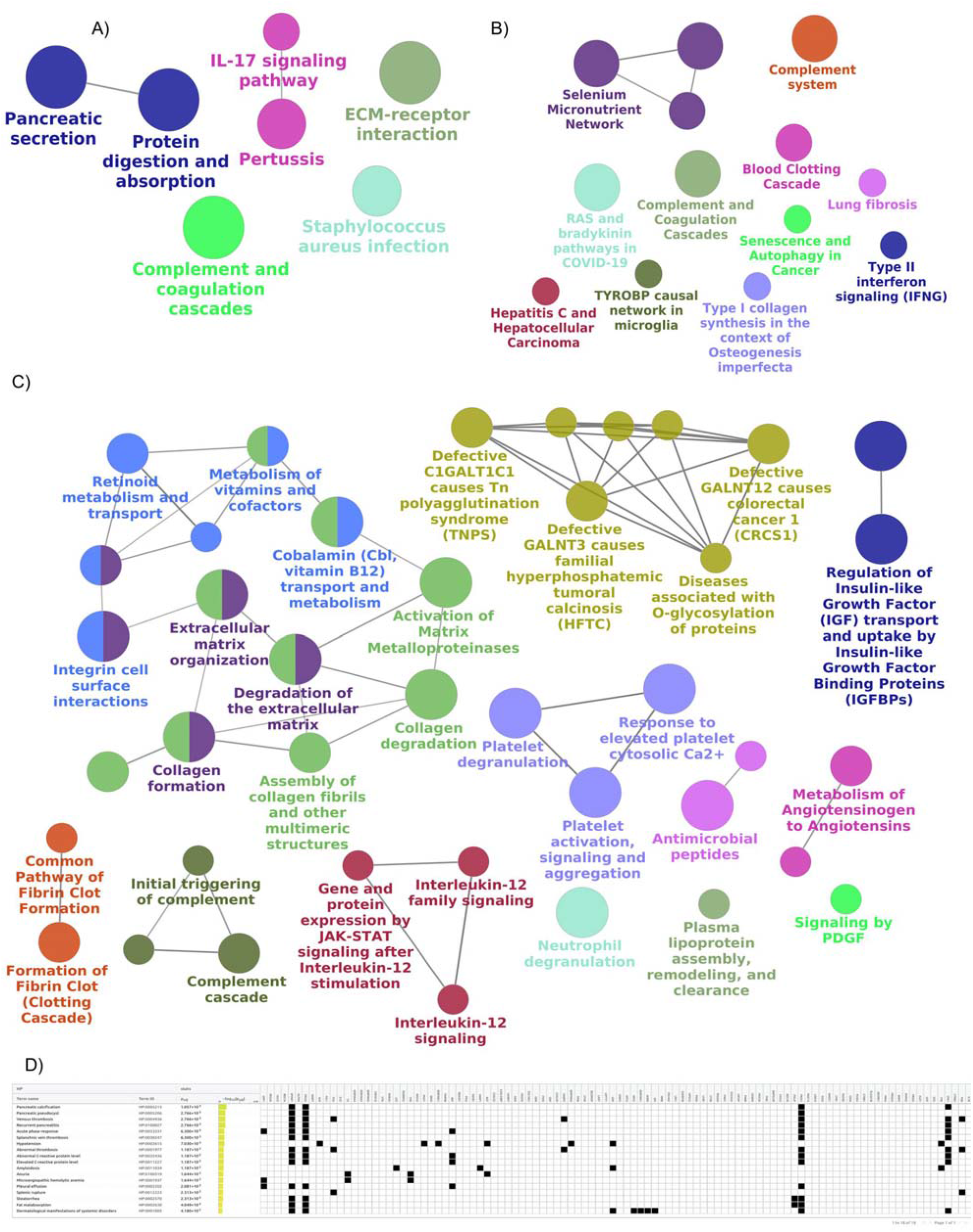
Enrichment analysis of biological pathways and disease phenome of the Secretome. A) KEGG pathways, B) WikiPathways, C) Reactome pathways, D) disease phenome from Human phenotype ontology.

Using the experimental evidence, we validated the mechanistic role of enriched biological pathways and their implications in MODs. The imbalance in Renin-Angiotensin system has been widely associated with Covid-19(Sriram et al., 2020). The RAS involves multiple organs and controls blood pressure, maintains fluid and electrolyte balance. The kidney secretes renin which acts on angiotensinogen (AGT), primarily secreted by the liver, to form angiotensin I(Ang I). We showed upregulation of *AGT* in SARS-CoV-2 infected pancreatic cells. Angiotensin-converting enzymes (ACE), present in the endothelial cells of the heart, lung, brain, kidney, convert Ang I to Ang II, which is vasoconstrictor and proinflammtory(Phillips and Kagiyama, 2002). It induces macrophage and IL-8 mediated neutrophil recruitment into the tissues through the endothelial lining of blood vessels(Liao et al., 2008; Nabah et al., 2004). AngII activates NADPH oxidase, which produces ROS. ROS leads to a reduction of NO and causes vasoconstriction. Ang II increases the production of a cytokine such as TNF-*α*, IL-1, and IL-6, CAM. The CAM stops immune cells and helps bring them into the tissues causing inflammation. It also prevents platelets, leads to coagulability and thrombus formation, and initiates atherosclerosis(Dandona et al., 2007; Dandona et al., 2003). Ang II mediates transcytosis of plasma low-density lipoprotein (LDL) particles across endothelial barriers, marking atherosclerosis’s start (Bian et al., 2017). In addition, Ang II plays a role in tissue fibrosis through angiotensin type 1 receptor (AT1) in cardiac, renal, pulmonary, abdominal tissues, and systemic sclerosis(Murphy et al., 2015). Pulmonary fibrosis impairs pulmonary function affecting the oxygen exchange in Covid-19 patients (Razzaque and Taguchi, 2003; Zou et al., 2021). Using pathway analysis, we demonstrate that pancreatic secretions are associated with RAS dysfunction, causing the immune system’s hyperactivation and coagulation abnormalities leading to multi-organ failure.

The disease phenotypes enrichment analysis of pancreatic Secretome using Human Phenotype Ontology revealed pancreatic calcification, pancreatic pseudocyst, venous thrombosis, recurrent pancreatitis, pleural effusion, splenic rupture, acute phase response, hypotension, abnormal thrombosis, elevated c-reactive protein level, amyloidosis, anuria, microangiopathic hemolytic anemia, fat malabsorption disease phenotypes (Fig.3 D). The documented experimental evidence indicates that pancreatitis leads to acute phase response, coagulation, thrombus formation, hemolytic anemia, amyloidosis. The amyloid deposition occurs in many organs such as the heart, kidneys, liver, spleen, nervous system, digestive tract. It induces inflammation, thrombosis, and immune dysfunction and causes systemic complications [86]. Thus, suggest the role of the upregulated pancreatic secretome-associated diseases phenotypes in MODs.

Interestingly, we found that FGB, FGG, ANXA2, MDK, AGT, VTN, SERPING1, CD44, IL1B are involved in many processes (Table S3). For example, blood coagulation: FGB, FGG, and SERPING1, vasoconstriction: AGT, pro-inflammatory response: IL1B, host-virus interaction: ANXA, cytokine and growth factor: MDK, cell adhesion and extracellular matrix organization: VTN and CD44. Furthermore, experimental evidence suggests that SARS-CoV-2 induced tissue damage, Renin-Angiotensin system (RAS) dysregulation, EC damage and thrombo-inflammation, Immune responses dysregulation, tissue fibrosis are fundamental processes of viral sepsis and MODs in COVID-19[57]. Therefore, our finding of upregulated pancreatic Secretome presents strong indications of the sepsis-mediated MODs.

### Gene-disease association network analysis of the pancreatic Secretome

We generated a gene-disease association network to further understand the implications of upregulated pancreatic Secretome in MODs (Fig.4 A). The top enriched disease classes were associated with Nervous, Cardiovascular, Metabolic, Immune, And Digestive Diseases (Fig.4 B), and suggesting multi-organ impact of upregulated pancreatic secretome. In addition, Our analysis revealed that IL1B, AGT, ALB, SPP1, CRP, SERPINA1, C3, TFRC, TNFSF10, and MIF proteins are associated with diverse disease terms in the gene-disease association network (Fig. 4 C-L). In addition, we found that IL1B, AGT, ALB, SPP1, CRP, SERPINA1, C3, TFRC, TNFSF10, and MIF were influential as they were linked to 31, 42, 47, 17, 24, 24, 17, 8, and 6 neighboring secretory proteins, respectively in the protein interaction network (Fig 2 B). We noted that IL1B was associated with 231 disease terms and 17 diseases classes, mainly with Nervous System and Cardiovascular Diseases. AGT was associated with 146 diseases and 15 classes, mainly with the Cardiovascular, Nervous System, and Digestive system. ALB was associated with 123 diseases and 17 classes, mainly with Urinogenital Disease and Pregnancy Complications, Immune System, Digestive System, and Cardiovascular Diseases. SPP1 was associate with 81 diseases and 13 classes, mainly with Nervous System, Digestive, Respiratory Tract Disease, and Cardiovascular Diseases. CRP was associated with 80 diseases and 16 classes, mainly associated with Cardiovascular, Digestive systems, Metabolic Disease, and Mental Disorders. SERPINA1 was associated with 59 diseases of 14 categories, primarily with Respiratory Tract and Digestive System Diseases. C3 was associated with 54 diseases of 13 types, mainly with Cardiovascular Disease and Nervous System Immune System Diseases. TFRC was associated with 52 diseases of 13 classes, mainly Hemic and Lymphatic Diseases and Immune System Diseases. TNFSF10 was associated with 49 diseases from eight different categories. Urogenital Diseases and Pregnancy Complications and Digestive System Diseases were among the top enriched disease classes. MIF is associated with 49 diseases of 11 classes. Skin and Connective Tissue Diseases, Mental Disorders, and Immune System Diseases were the top enriched disease classes. Thus, our analysis suggests that upregulation of *IL1B, AGT, ALB, SPP1, CRP, SERPINA1, C3, TFRC, TNFSF10*, and *MIF* genes may have systemic effects and may impact MODs.

**Fig. 4.**
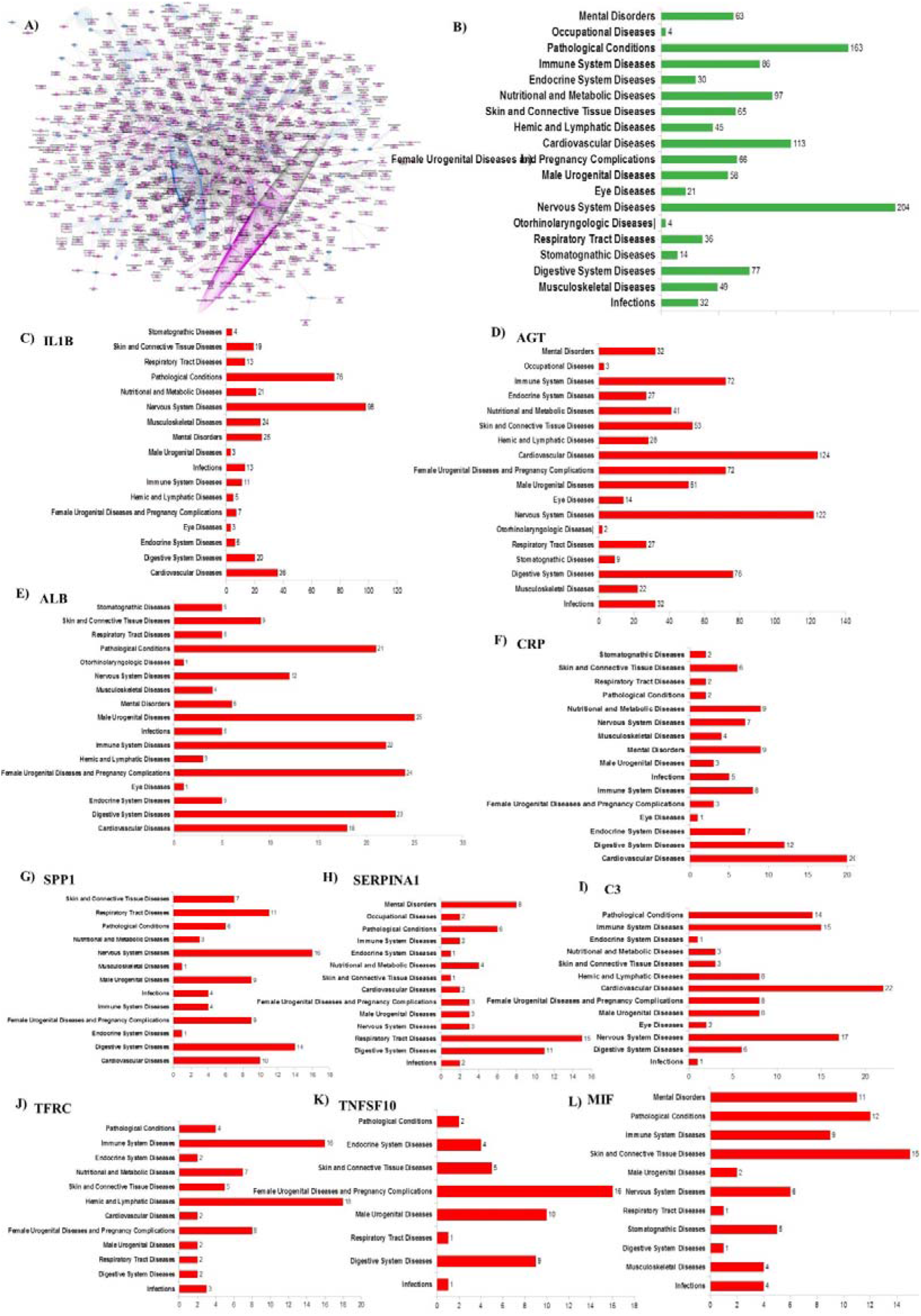
A) The Gene-Disease association network of secretome genes. Genes and diseases are represented as blue and pink nodes. Bar plot showing enriched disease classes for B) Secretome, C) IL1B, D) AGT, E) ALB, F) SPP1, G) CRP, H) SERPINA1, I) C3, J) TFRC, K) TNFSF10 and L) MIF

## CONCLUSION

Using the scRNA-seq data analysis of SARS-CoV-2 infected pancreatic cells, we identified the upregulated pancreatic Secretome and linked it with the COVID-19 associated multi-organ dysfunction. We validated, using extensive literature and experimental evidence, that upregulated pancreatic Secretome is associated with coagulation cascade, complement activation, renin angiotensinogen system dysregulation, endothelial cell injury and thrombosis, immune system dysregulation, and fibrosis. In addition, we showed that the upregulated and influential *genes IL1B, AGT, ALB, SPP1, CRP, SERPINA1, C3, TFRC, TNFSF10*, and *MIF* may have systemic effects and lead to multi-organ dysfunction. Our analysis suggests the influence of upregulated Secretome on multi-organ systems such as Nervous, Cardiovascular, Immune, Digestive, and Urogenital systems. Thus, our results indicate that the dysregulation of the pancreatic Secretome has a multi-organ impact in COVID-19. Future progress needs to understand the triggers involved in upregulating the pancreatic Secretome to curb the associated multi-organ damage.

## Author contributions statement

E.P conceived and designed the research; E.P performed literature survey, Single cell RNAseq data analysis and prepared the illustrations; E.P, and RM analyzed the data; E.P and R.M wrote the manuscript. R.M supervised the whole study. All the authors approved the final version of the manuscript before submission.

## Acknowledgments

Computational facility support to R.M from the Banaras Hindu University is also gratefully acknowledged.

## METHODS

scRNA-seq data of ex vivo SARS-CoV-2 infected human pancreas were retrieved from Gene Expression Omnibus(GSE159556)(Tang et al., 2021). Two samples each for Mock-infected and SARS-CoV-2 infected were taken. scRNA-seq data analysis was performed using Seurat 4.0.2.(Hao et al., 2021)

### Single-cell RNA-seq data analysis

#### Quality control

We filtered cells having less than 200 genes, greater than 20% mitochondria gene, and less than 5% ribosomal genes. In addition, the cells having genes expressed in less than 3 cells are also filtered. Then we removed the effects of the cell cycle on the transcriptome by using CellCycleScoring. Before running CellCycleScoring, the data was normalized and logtransformed using NormalizeData. We then removed the doublets using DoubletFinder(McGinnis et al., 2019). The doublet prediction was run on each sample separately with a 4-7.6% doublet rate based on the loading rate. After removing doublets, the two SARS-CoV-2 infected samples now have 5821 and 6661 cells, whereas the two Mock-infected samples have 3726 and 7869 cells. Next, we used the QC-filtered data to identify the top 2000 variable genes using FindVariableFeatures with selection.method “vst”. Next, ScaleData was used to scale and center the data, where the number of genes and percentage of mitochondrial genes were the “vars.to.regress”. After scaling, we performed principal component analysis (PCA) and Uniform Manifold Approximation and Projection(UMAP) for dimensionality reduction using the first 30 principal components.

#### Integration and clustering

We then used FindIntegrationAnchors to identify anchors in Seurat objects and integrated the datasets with IntegrateData(). Then, the dataset was scaled using ScaleData. The PCA and UMAP were performed using the first 30 dimensions. Next, we used FindNeighbors to Compute the nearest neighbor graph using the top 30 PCs. We then performed the graph-based clustering using FindClusters at a resolution of 4.5. Clustree package was used to choose the final resolution (Zappia and Oshlack, 2018).

#### Cell type identification

We first identified genes differentially expressed in a cluster with respect to other clusters using FindAllMarkers with logfc.threshold = 0.25, min.pct = 0.25, min.diff.pct = 0.25. Benjamini-Hochberg false discovery rate (FDR) was 0.05. The test used was Wilcoxon Rank Sum test, and the assay was “RNA”. We used literature to manually curate DEGs of each cluster to identify cell types. After identifying the cell type for clusters, we merged the same cell type cluster into one. This resulted in nine clusters of acinar cells, ductal cells, alpha cells, beta cells, delta cells, PP cells, endothelial cells, mesenchyme cells, and immune cells.

#### DEGs across SARS-CoV-2 infected and Mock-infected conditions

We have identified differentially expressed genes between SARS-CoV-2 infected and Mock-infected conditions using FindAllMarkers with logfc.threshold = 0.2, min.pct = 0.1 at FDR =0.05. We used the Wilcoxon Rank Sum test on “RNA” assay.

#### Identification of secretome

We identified the secretory proteins using Human Protein Atlas (Uhlen et al., 2017). Then, utilizing this protein set as input, we regenerated the Protein interaction network using STRING(Szklarczyk et al., 2019). Finally, we used Cytoscape 3.8.2 to visualize and analyze the network (Paul et al., 2003).

#### Enrichment analysis

We used g:Profiler for GO enrichment analysis, Biological pathway enrichment analysis using KEGG, Reactome, WikiPathways. The disease phenotypes enrichment analysis was performed using Human Phenotype Ontology)(Raudvere et al., 2019) ClueGO, a Cytoscape plug-in, was used to identify the functionally grouped GO and pathway(Bindea et al., 2009). We used the DisGeNET Cytoscape app (7.3.0) for gene-disease associations (GDAs)(Piñero et al., 2016). We used EnhancedVolcano for generating a volcano plot (Blighe et al., 2019). Pheatmap package was used for generating heatmap(Kolde and Kolde, 2015).

